# Disproportionate feedback interactions govern cell-type specific proliferation in mammalian cells

**DOI:** 10.1101/323147

**Authors:** Dola Sengupta, Vijay Phanindra Srikanth Kompella, Sandip Kar

## Abstract

In mammalian cells, the critical decision to maintain quiescence over proliferation commitment in and around the G_1_-S transition depends on more than one intertwined feedback interaction, and is highly cell-type dependent. However, the precise role played by these individual feedback regulations, in order to generate such diverse nature of proliferation commitment, is still poorly understood. Herein, we propose a generic mathematical model of G_1_-S transition in mammalian cells that not only reconciles distinct single cell experimental observations in a cell-type specific manner, but also makes experimentally testable non-intuitive predictions. Intriguingly, The model analysis reveals that the feedback motifs responsible for the G_1_-S transition act in a disparate fashion to organize the cell-type specific proliferation response in different mammalian cells. The proposed model, in principle, can be effectively tuned to explore proliferation dynamics in a cell-type specific way to gain crucial insights about novel therapeutic intervention to prevent unwanted cellular proliferation.

## Introduction

Commitment to cellular proliferation from a quiescent (G_0_ or G_1_ like state) phase of the cell cycle is one of the most tightly regulated processes in any eukaryotic organism. In mammals, only few cell types are actively undergoing growth factor (GF) mediated cell cycle in a regular interval, but in a controlled manner^1–5^. Most of the other cells in mammals, in spite of being experiencing similar GF condition, remain in a quiescent phase^6^. Depending on the need, even these quiescent cells for a certain higher threshold level of GF can eventually commit for active proliferation cycle by successfully making the G_1_-S transition^7^. In many cancerous cells, the G_1_-S transition gets deregulated in one way or other, leading to uncontrolled cellular proliferation commitment even at much lower GF condition^8^. Importantly, recent single cell experimental studies with different mammalian cells have uncovered that the events regulating G_1_-S transition were highly cell-type dependent^7,9,10^. It is therefore imperative to understand the dynamics of the G_1_-S transition in mammalian cells preferably in a cell-type specific fashion to invent novel preventive measures to inhibit uncontrolled proliferation at the early stage of the cell cycle.

It was well known in the literature that the family of E2F transcription factors (especially E2F1) plays a crucial role in triggering the proliferation response by initiating the G_1_-S transition^7,8,11–14^ On the other hand, Rb (Retinoblastoma protein) prevents the progression towards proliferation by inhibiting E2F^7,11^. Lately, Yao et al. had convincingly demonstrated that the Rb-E2F network functions as a bi-stable switch in mouse fibroblast cells (REF52 cells) to transform a graded growth factor stimulus into all or none E2F response, and dictates decision to undergo cellular proliferation or quiescence in the late G_1_ phase of the cell cycle^7^. Intriguingly, several cell cycle regulatory proteins through multiple feedback (positive, negative and feed-forward) interactions, control the E2F activation dynamics within a mammalian cell. Thus, one can speculate that in different mammalian cells, subtle variations in these feedback interactions might lead to the cell-type specific dynamics of G_1_-S transition.

Indeed, the disparity in these feedback interactions is evident in different cell types. In mouse fibroblast cells (REF52 cells) it had been observed that Myc protein mainly controls the amplitude of E2F activation^9^. Interestingly, CDK’s (Cyclin dependent kinase) like CDK4 and CDK2 (aided by Cyclin D and Cyclin E, respectively) had little effect on E2F amplitude, but the duration of the E2F activation was determined by these CDK’s^9^. Whereas, in MCF10A cell line, an immediate build up of CDK2 activity after mitosis was shown to be essential for the proliferation onset, and the cells transit into a G_0_-like state if the CDK2 activity is suppressed in the new born cell^15,16^. The CDK2 activity in MCF10A cells critically depends on the level of the CKI’s (Cyclin dependent kinase inhibitor) like p21 (CDK2 inhibitor) at the end of the previous cell cycle^10^. In the similar note, Barr et al. showed that in Hela cells, a switch like G_1_ to S phase entry is mainly driven by the double-negative feedback loop between Cyclin E/CDK2 and CDK inhibitor p27^17^. Recently, Cappell et al. had revealed that reaching certain amplitude of E2F does not ensure proliferation commitment^16^. They showed that the point of no return for a cell towards quiescence is confirmed by a rapid APC^Cdh1^ inactivation via Cyclin E/CDK2 to trigger the G_1_-S transition^16^. All these live cell-imaging studies importantly emphasize the fact that the proliferation decision-making events are extremely cell-type specific, and no wonder regulated by a set of complicated events and feedback motifs.

In literature, mathematical and computational modeling had been used quite often and successfully to understand the complicated mammalian cell cycle dynamics, especially in the context of G_1_-S transition^18–23^. Even in the latest single cell experimental studies, there were attempts to supplement the experimental data by constructing phenomenological mathematical models^7,9,10^. These models^7,9,10^ partly describe some of the observations made in the experiments, but often fail to capture many key aspects of the experimental findings in a cell-type specific way, and lack predictive power. This can be due to many reasons. Firstly, most of these models are built by using phenomenological terms (like Hill or Michaelis-Menten kinetics) without considering the actual molecular mechanism behind such complex regulations. Second, in many of these models the inherent molecular fluctuations were ignored while developing the model. Most importantly, these models are built in such a way that it is very difficult to convert a particular model developed for a specific cell-type to another model corresponding to a different mammalian cell.

Keeping these requirements in mind, in this article, we have constructed a mass-action kinetics based predictive mathematical and computational model of G_1_-S transition for mammalian cells to elucidate the single cell experimental data in a cell-type specific fashion. Our model includes experimentally observed basic molecular interactions in terms of mass-action kinetics, which makes it suitable for stochastic analysis by employing Gillespie stochastic simulation algorithm (SSA)^24^ We have demonstrated that the model can be intuitively tuned to corroborate single cell experimental observations for different cell-types, and at the same time is capable of making novel predictions for the particular cell type concerned, which are feasible experimentally. Importantly, our modeling study reveals how different mammalian cells distribute the labor for organizing the G_1_-S transition among a set of feedback motifs to generate diversity in the proliferation commitment. Thus, we believe that our generic model can be effectively modified to disentangle the proliferation dynamics of mammalian cells in a case-by-case basis.

## Mathematical model of G_1_-S transition in mammalian cell

The model depicted in Fig. 1 (detail description is given in Fig. S1 and SI Text) describes how E2F dynamics is controlled in a mammalian cell by various activator and inhibitors of the cell cycle control machinery during the R-point as well as G_1_-S transition in response to growth factor stimulation. It is well known in literature that in quiescent or non-cycling cells, Rb represses E2F activation by sequestering E2F making a hetero-dimeric complex^7,8,12–14^. Upon growth factor stimulation, transcription factor Myc activates the production of CycD, which in turn forms the CycD/CDK4 complex that phosphorylates Rb resulting in the partial release of E2F from Rb-E2F complex^7–9,11,12,25,26^. The free form of E2F then gets involved into one direct and another indirect positive feedback loops to activate its own transcription and activation. Firstly, the free E2F forms a hetero-dimeric complex (DE) with the Dimerization partner protein (DP), which ultimately acts as transcriptional activator of E2F itself, and for other proteins such as Myc, CycE etc.^7,8,12–14^.

**Fig. 1.**
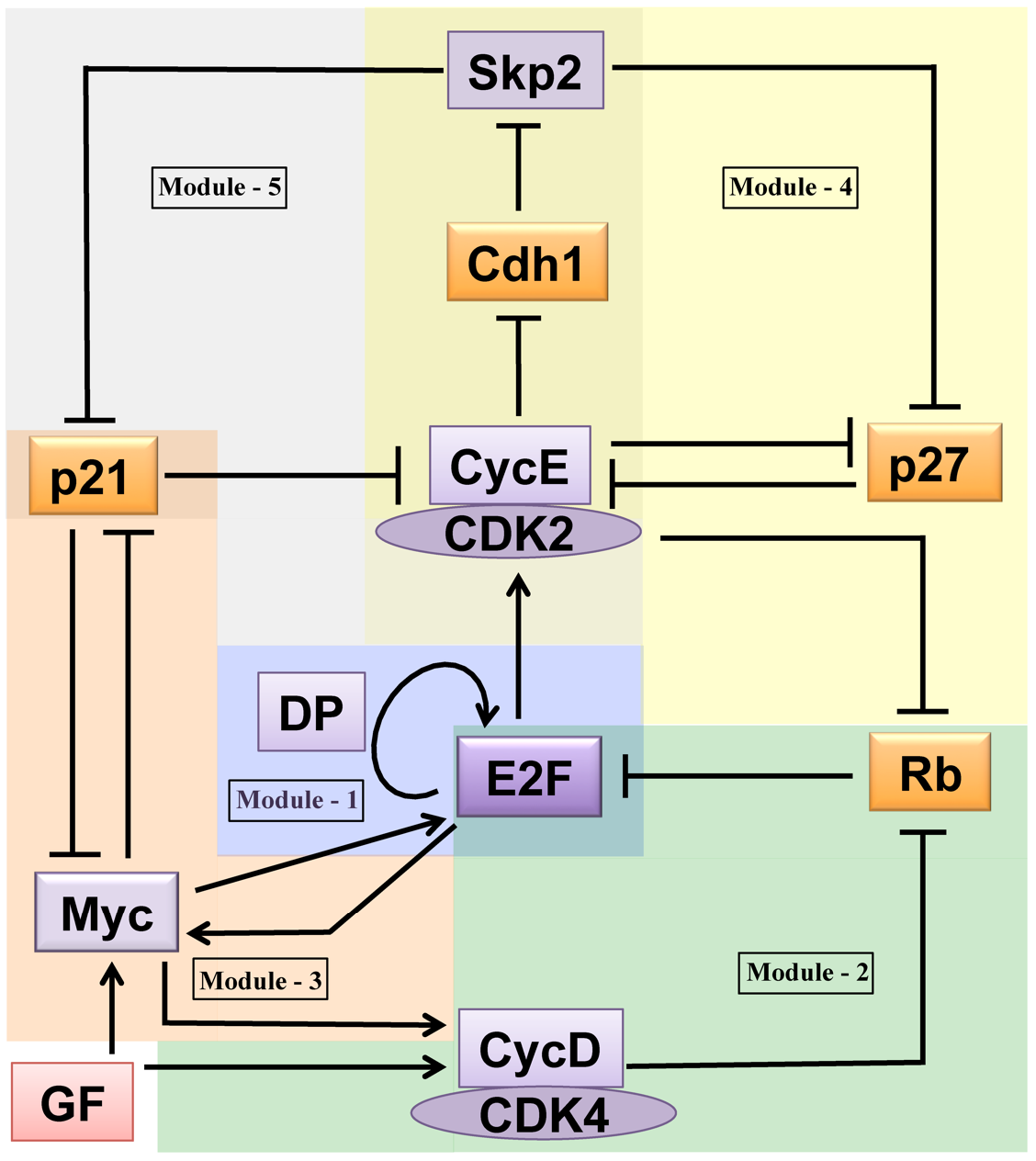
Proposed network for the GF-driven G_1_-S transition model in Mammalian cell. (Solid arrows represent activation and hammer headed lines represent inhibition, respectively.) The overall network can be divided into 5 connected modules. **Module-1** represents how E2F positively regulates its own transcription; **Module-2** depicts how GF mediated CycD activation modulates the Rb-E2F interaction; **Module-3** reflects how GF regulates the mutual antagonistic interaction between Myc and p21; **Module-4** and **Module-5** describe how E2F promotes CycE-Cdk2 activity by influencing the mutual antagonistic interactions between CycE-Cdk2 and CKI’s like p27 (in module-4) and p21 (in module-5) by promoting Skp2 mediated p27 and p21 degradation. Fig. S1 provides a more detailed description of all the interactions considered in these 5 modules, and the corresponding kinetic equations, description of the variables and parameters are depicted in Tables S1-S5, respectively.

Once produced, CycE forms a hetero-dimeric complex with CDK2 (CycEC) and initializes the hyper-phosphorylation of Rb leading to complete removal of Rb repression over E2F^10^. This inactivation of Rb although is not a straight forward event due to the fact that the CycE/CDK2 complex after its production immediately gets sequestered in trimeric-complex with the CKI inhibitors like p21 and p27^17,26–28^. Once there is enough accumulation of the CycE/CDK2 in the system it inactivates Cdh1 that in turn helps in the release of Skp2^10,15–17,27–30^. Skp2 recognizes potent Cyclin-CDK inhibitors p21 and p27 bound with CycE/CDK2 complex, and initiates the ubiquitination of the phosphorylated forms of CKI’s by enhancing their proteasomal degradation^10,15–17,29–32^. CycE/CDK2 complex further facilitates the degradation of p27 by phosphorylating it (by phosphorylating p27 at T187 targeting p27^Kip1^ to the proteasomal machinery)^10,17,31,32^. On the other hand, literature suggests that the potent oncogene Myc orchestrates E2F activation even during the very early stage of the G1 phase by being involved in a double negative feedback as well as in a positive feedback loop interactions with p21 and E2F, respectively^8,9,33–37^.

The overall model described in Fig. 1 is converted into mass-action kinetics based (with only one phenomenological term) ordinary differential equations (Table S1). We have given a detail description of the proposed mathematical model in SI Text by defining the variables involved in Table S2. The definitions of the model parameters along with their numerical values are illustrated in Tables S3-S5. It is important to mention that here our endeavor is to capture the complex dynamic behavior of E2F in cell-type specific manner by analyzing a generic regulatory network that controls the R-point as well as the G_1_-S transition in different mammalian cells as a function of growth factor. In that venture, we have chosen most of the parameters involved in the model in an intuitive way with a goal to cultivate the effect of different feedback loops present in the network while maintaining reasonable expression levels of different network components.

## Results

### E2F steady state shows tri-stability as a function of growth factor in REF52 cells

The bifurcation analysis of the proposed model (Fig. 1 and Fig. S1, Tables S1-S3) indicates that under certain parametric situation the E2F steady state level can essentially function as a tri-stable switch (Fig. 2a) with increasing doses of growth factor (GF). In Fig. 2a, the stable region (indicated by red line) in the E2F steady state (between the saddle nodes SN_1_ (at 0.34%) and SN_2_ (at 0.04%)) that appears for relatively low growth factor stimulation delineates the existence of a quiescent G_0_ like state during the cell cycle progression. At the same time, the E2F steady state region between the saddle nodes SN_2_ (at 0.04%) and SN_3_ (at 1.84%) denotes the G_1_ like state of the cell cycle (since E2F still remains in an ‘OFF state’ as number of E2F molecules are low) while, the upper part of the E2F steady state bifurcation diagram (starting from the saddle node SN_4_ (at 0.003%)) signifies the S-G_2_-M like phase of the cell cycle (where E2F remains in an ‘ON state’ as number of molecules is relatively high). This tri-stability in E2F steady state dynamics as a function of GF suggests that initially with increase in GF, the quiescent cells from a G_0_ like state can transit to a G_1_ like state. Once the GF crosses a certain critical threshold, the cells residing in the G_1_ phase can further commit to the S-G_2_-M (ON state) phase of the cell cycle.

**Fig. 2.**
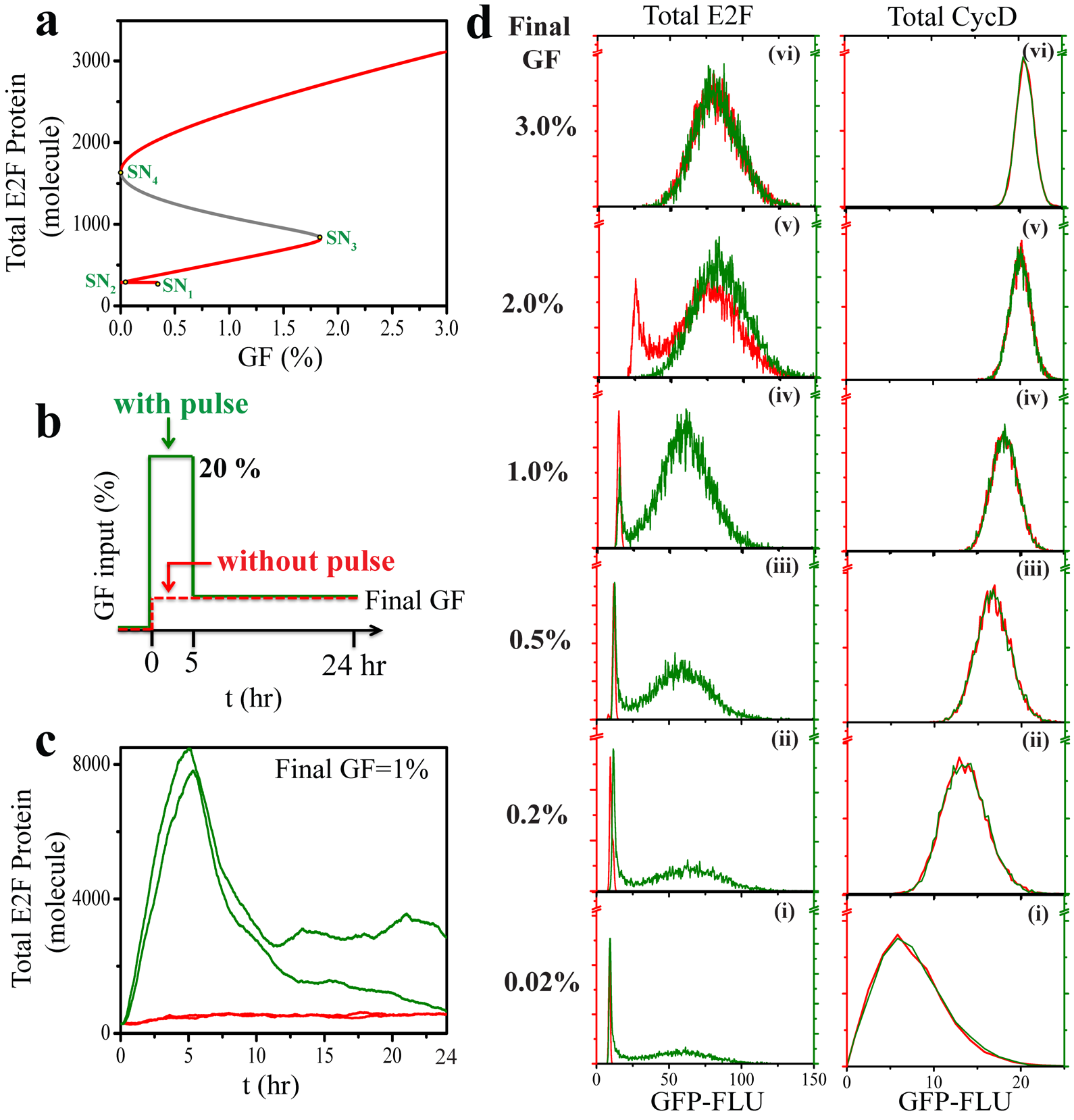
A tri-stable E2F steady state dynamics explains the history dependence behavior of E2F in REF52 cells. (**a**) The E2F steady state dynamics as a function of GF shows tri-stability. (**b**) The protocol followed for numerical simulation to mimic GF stimulation experiments performed with serum starved cells under pulsed (20% for 5 hrs, green line) and non-pulsed (red line) conditions. (**c**) Few representative stochastic trajectories under pulsed (green line) and non-pulsed (red line) conditions, where the final GF is set to 1%. (**d**) Histograms of total E2F and Total CycD levels are plotted for 10000 cells (i.e., in our case for 10000 different starting random number seeds during stochastic simulation) after 24 hrs of GF stimulation for pulsed (green line) and non-pulsed (red line) situations. Final GF percentages are set to (**i**) 0.02% (**ii**) 0.2% (**iii**) 0.5%, (**iv**) 1%, (**v**) 2% and (**vi**) 3%, respectively. To compare the histograms with Yao et al.’s FACS experiment^7^, Fig. S2 shows the histograms in terms of number of molecules of E2F and CycD. (For details see method section, parameters are given in Table S3)

We envisage that a similar E2F steady state dynamics is operative in the REF52 cell-type. To justify our conjecture, we performed a set of stochastic simulations to reconcile the history-dependence of E2F activation in these REF52 cells as observed experimentally by Yao et al.^7^. History-dependence marks the characteristic property of a system whose output responds with a delay to the change in the input, or does not totally regain its original state. To examine that Yao et al.^7^ induced the output to an admissibly high steady state level of E2F with a strong input (20% GF for 5 hours), and then finally set the input to a lower level to inspect the new output steady state behavior (as shown in Fig. 2b) via FACS measurement technique considering around 10000 cells. In the similar fashion we have performed our stochastic simulation for 10000 cells (simulating 10000 different stochastic trajectories by changing the random number seed during SSA simulation, few representative trajectories are shown in Fig. 2c) and recorded the E2F responses at a range of growth factor input (0.02-3% GF for 24 hour) with or without an initial sufficiently strong growth factor pulse (20% GF for 5 hours) (Fig. 2d).

Fig. 2d reveals that at final GF concentrations corresponding to 0.02% to 1%, cells are residing in the ‘OFF state’ with low E2F response in the numerically simulated non-pulsed situation (Fig. 2d (left column, (i-iv)), red lines). On the contrary, in the case of stochastic simulations performed with a GF pulse (20% GF for 5 hours, Fig. 2d (left column, (i-iv)), green lines), cells mostly reside in the ‘ON state’ with high E2F level. Interestingly, a numerical pulse (20% GF for 5 hours, Fig. 2d (left column), green lines) experiment leads to bimodal E2F distribution within a cellular population even when the final growth factor concentration is set as low as 0.02% or 0.2% (i-ii), which is in very close resemblance with the experimental observation made by Yao et al.^7^ with REF52 cells. For the non-pulsed case, our stochastic simulation suggests that bi-modality in E2F distribution will be observed in between 1% to 2% GF condition that further corroborates with the experimental findings^7^. In comparison to the E2F response, the changes in the CycD level in our stochastic simulations for the same population of cells happen in a uni-modal manner irrespective of the pulsed or non-pulsed conditions (Fig. 2d (right column)). Thus, not only our model simulations qualitatively substantiate the history dependence experiments performed by Yao et al. with REF52 cells^7^, but also it can further capture even finer details like the slight shift of the peaks in the E2F (as well as CycD) responses in case of both non-pulsed and pulsed cases (0.02-1% GF, Fig. S2) as observed by Yao et al.^7^. This can be attributed to the presence of a G_0_ like state in the E2F steady state dynamics for the lower level GF as predicted by the deterministic bifurcation diagram of E2F versus GF (Fig. 2a). We observe such a shift from a G_0_ like to G_1_ like state in our stochastic simulations (shown in Fig. S2(i-iv), inset), and this kind of strengthens our prediction from the bifurcation analysis (Fig. 2a) that with respect to E2F level as a function of GF, a closely separated G_0_ and G_1_ like states exist in REF52 cells.

### Cyclins control the time required to attain E2F ON state but Myc modulates the amplitude

At this juncture, we asked whether our model could account for other important experimental observations made for REF52 cells or not. In this regard, Dong et al. had recently shown that threshold amplitude of E2F in individual cells is a reliable predictor of the cell cycle commitment by taking REF52 cells as the reference system^9^. Additionally, by using CDK4/2 inhibitors, Dong et al. demonstrated that CycD/E activities do not affect the E2F amplitude in REF52 cells^9^. Our bifurcation analysis indicates that indeed CycD and CycE both individually (Fig. S3a, presence of CDK4 inhibitor and Fig. S3b, presence of CDK2 inhibitor) and collectively (Fig. 3a(i), left panel, presence of both the inhibitors) hardly affect the E2F steady state levels as a function of GF. To mimic the effect of the corresponding inhibitors in the model, we have reduced the transcription rates of CDK4/2 keeping other model parameters constant as in Table S3. Stochastic simulation at 3% GF for the individual and dual CDK4/2 inhibitor conditions show almost no deviation of the E2F steady state distribution (Fig. S3a-b, lower panels and Fig. 3a, right panel) in comparison to the WT situation (Fig. S2(vi), left panel (non-pulsed case)).

**Fig. 3.**
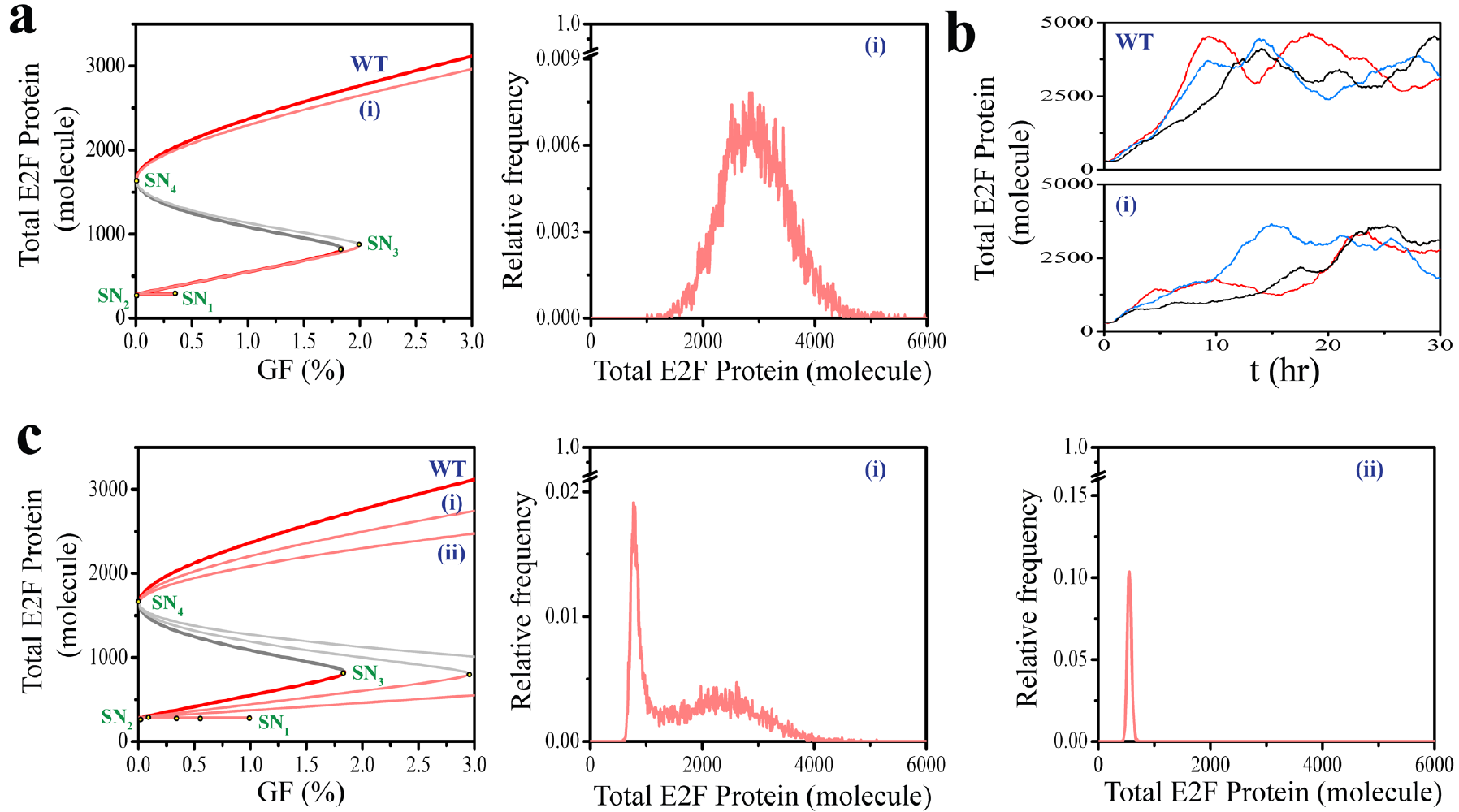
Cyclins control the time required to attain E2F ON state, but Myc modulates the amplitude of E2F in REF52 cells. **(a)** The E2F steady state dynamics as a function of GF under wild type (WT) and both CycD and CycE inhibitory **(i)** conditions (**left Panel**, *k*_mK4_=4 molecule min^−1^, *k*_mK2_=0.4 molecule min^−1^). The histogram (following same protocol as in Fig. 2) of E2F level for 3% GF (non-pulsed) and under simultaneous CycD and CycE **(i)** inhibition shows no significant change in amplitude of E2F distribution (**right panel**) in comparison to Fig. 2d(vi) (or Fig. S2d(vi)). **(b)** Few representative stochastic trajectories of E2F activation under 3% GF (non-pulsed) condition reveal that time required to attain E2F ON state in WT (**upper panel**) is considerably less in comparison to the simultaneous CycD and CycE **(i)** inhibitory condition (**lower panel**, *k*_mK4_=4 molecule min^−1^, *k*_mK2_=0.4 molecule min^−1^). **(c)** The E2F steady state dynamics as a function of GF under wild type (WT) and for moderate **(i)** (*k*_y9_=0.288 min^−1^) and high **(ii)** (*k*_y9_=0.16 min^−1^) levels of Myc inhibition (**left Panel**). The histograms (following same protocol as in Fig.2) of E2F level for 3% GF (non-pulsed) under moderate (**i**, **middle panel**) and high (**ii**, **right panel**) levels of Myc inhibition display significant change in the amplitude of E2F distribution in comparison to Fig. 2d(vi) (or Fig. S2d(vi)). (Rest of the parameters is same as in Table S3)

This clearly implies that inhibition of CycD and CycE activities will have no effect in controlling the E2F amplitude for the REF52 kind of cells. At the same time, if we look at how E2F gets accumulated during the course of G_1_-S transition in few representative single cells corresponding to WT (Fig. 3b, upper panel) and under CDK4/2 inhibitory (Fig. 3b, lower panel) conditions, we clearly observe that in absence of cyclin D/E activities, there is a delay in the E2F activation. Thus, our model verifies that cyclin D and cyclin E modulate the duration of E2F activation in REF52 cells but have little or no effect on the E2F amplitude.

Dong et al. further reported that the transcription factor Myc, critically modulates the threshold amplitude of E2F in REF52 cells^9^. By employing different concentrations of inhibitor of Myc, they showed that the cell cycle entry could be altered via precisely modulating the E2F amplitude^9^. In our model, we simulated the inhibition of Myc by reducing its transcription rate. Our bifurcation analysis (Fig. 3c, left panel) predicts that increasing the extent of Myc inhibition results into shift of SN3 saddle node towards higher GF condition. This implies that if Myc is inhibited then the cells will not be able to transit to the S-G_2_-M like state at a relatively lower GF condition, which is possible for the WT scenario. Our stochastic simulations (Fig. 3c, middle and right panel), where we simulated the increasing Myc inhibition conditions, reveal that these inhibitory situations will lead to either a bi-modal (Fig. 3c, middle panel) or a uni-modal (Fig. 3c, right panel) E2F steady state distribution representing either a partially or fully quiescent population of REF52 cells for 3% GF condition. Importantly, at the same GF condition, the majority of the cells remain in a S-G_2_-M like state in the WT case (Fig. S2(vi), left panel, (non-pulsed case)). These results clearly show that our G_1_-S transition model can qualitatively substantiate the observations made at the single cell level for the REF52 cell types^9^.

### Model predicts the outcome of several new experiments related to REF52 cell-type

To test our proposed model further, we performed some numerical experiments that are in principle possible to validate by performing real experiments. We first tested the fate of the cells, if the pulse duration employed by Yao et al.^7^ in their pulsed experiments (with 20% GF) varies from 5 hr to a higher (6 hr) or a lower (4 hr) value. We performed this numerical experiment at GF = 0.3% without any pulse (Fig. 4a), with 4 hr (Fig. 4b) and 6 hr (Fig. 4c) pulse, and analyzed the level of E2F for about 10000 cells at the end of 24 hr in each case. A careful analysis of the Fig. 4a-c reveals that increase in the pulse duration definitely elevates the number of committed cells in the S-G_2_-M like state. Interestingly, Fig. 4a predicts that even if the cells are kept at 0.3% of GF medium after the GF starvation, a high proportion of cells will reside in a quiescent G_0_ like phase (Fig. 4a, inset) due to the existence of a low lying E2F (Fig. 2a) steady state under such a GF condition.

**Fig. 4.**
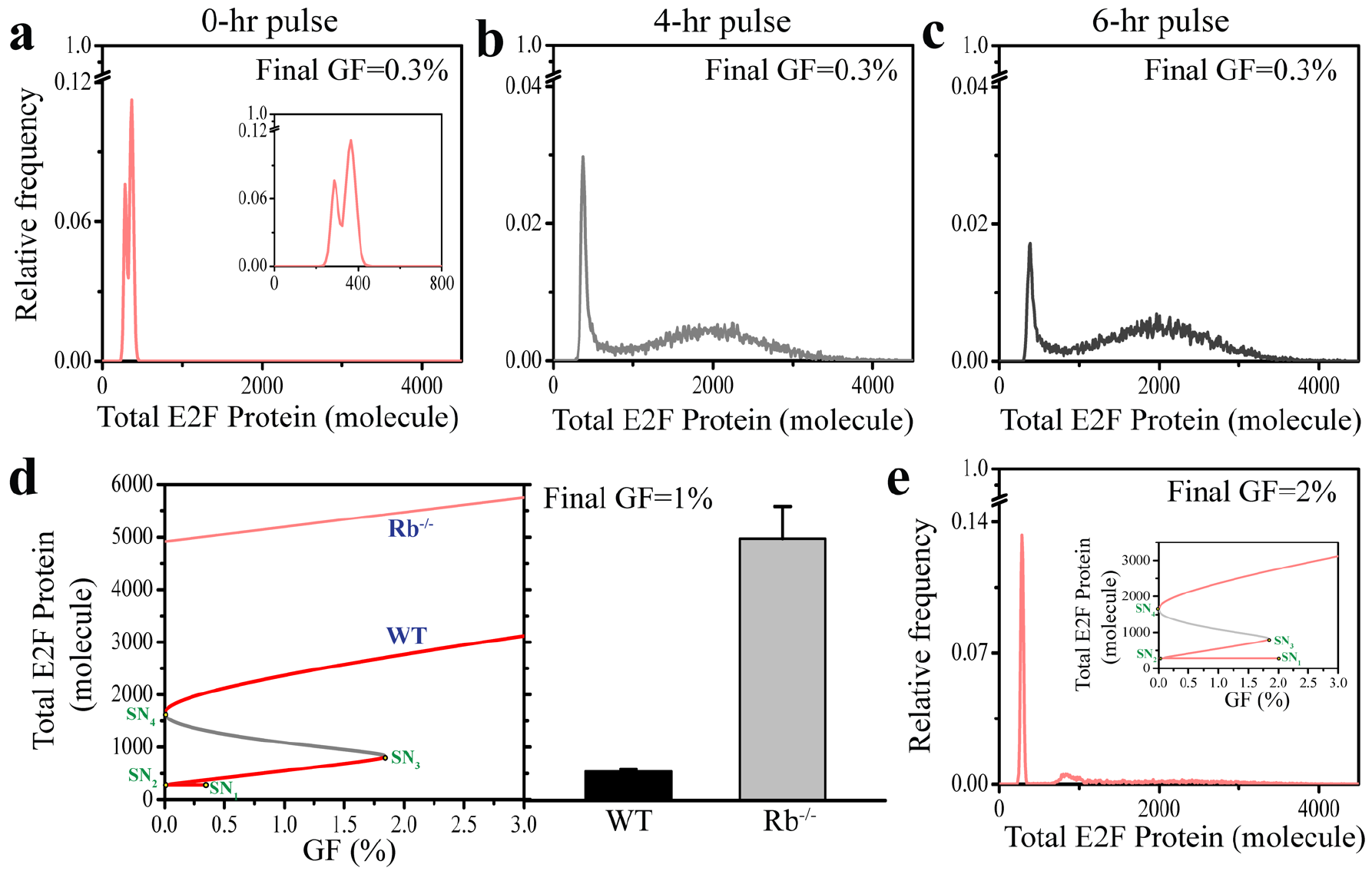
Model makes experimentally testable predictions for REF52 cells. The histograms of E2F level after 24 hrs of stochastic simulation (considering 10000 cells, following same protocol as in Fig. 2) with 0.3% GF (final level) under **(a)** 0-hr initial pulse (with two distinct peaks reflecting the simultaneous existence of G_0_ and G_1_ like states, **inset**), **(b)** 4-hr initial pulse of 20% GF, and **(c)** 6-hr initial pulse of 20% GF, respectively, reflect that the proportions of cells with E2F ON state increases significantly as the pulse duration is increased. **(d)** The E2F steady state dynamics as a function of GF under Rb^-/-^ condition (*k*_rnrb_=0 molecule min^−1^) has only one stable steady state with very high level of E2F in comparison to tri-stable wild type (WT) scenario, and even GF as low as 1% leads to G_1_ to S-G_2_-M like transition (under Rb^-/-^ condition). **(e)** The histogram (following same protocol as in Fig. 2) of E2F level for 2% GF (non-pulsed) in presence of silibinin (*k*_p21d_=0.2 X WT, *k*_npx_=0.2 X WT), displays almost a near uni-modal E2F distribution (with low level of E2F) that complies with the E2F steady state bifurcation diagram (**inset**) as a function of GF with an extended G_0_ like state, and significantly differs with the E2F distribution observed under WT situation (Fig. 2d(v) and Fig. S2d(v)). (Rest of the parameters is same as Table S3)

While, a 20% GF pulse of different higher durations allow more cells to stay in a S-G_2_-M like state (Fig. 4b-c). These predictions corroborates with the recent experimental study performed by Kwon et al. for the REF52 cell-type^38^, where they had seen similar increase in the proportion of committed cells with increased pulse duration.

The other kinds of experiments one can perform are either (i) knocking-out a gene, or (ii) applying an inhibitor related to a particular gene, or (iii) overexpressing a particular gene related to the network. Keeping this in mind, we executed some numerical experiment in these directions as well. Since the Rb-E2F regulation plays a major in regulating the G_1_-S transition in REF52 cells, we first performed a bifurcation analysis for a Rb knockout situation (Fig. 4d). Fig. 4d reveals that under such a condition, the REF52 cells will commit to S-G_2_-M like state even for very low growth factor level (say GF = 1%), which will lead to uncontrolled proliferation. Whereas, the deterministic bifurcation analysis (Fig. 4e, inset) portrays that if silibinin (stabilizes p21 and p27 proteins) is employed in the cell culture medium, then the cells will have an extended G_0_ like state. Thus, even for a relatively higher GF condition (GF = 2%) quite a high fraction of cells will remain in a quiescent phase of the cell cycle (Fig. 4e). We further performed a systematic sensitivity analysis (Fig. S4a-b) on the mRNA transcription rates for all the components involved in the G_1_-S transition network to know how overexpressing any of those mRNA’s are going to alter the dynamics significantly. The analysis predicts that overexpressing four different mRNA’s (namely DP (Fig. S4c), p21 (Fig. S4d), Skp2 (Fig. S4e) and Cdh1 (Fig. S4f)) will significantly alter the G_1_-S transition dynamics in case of REF52 cells.

### p21 regulates growth factor-dependent population heterogeneity in MCF10A cells

It was evident from the analysis of the proposed model that under the current parametric condition, it mimics qualitatively the experimental observations for REF52 cells. Can this model (Fig. 1, Fig. S1 and Table S1) be utilized as a general framework to model G_1_-S transition for other cell-types? To explore this issue, we focus on the recent experimental works by Overton et al. and Cappell et al. on MCF10A cell line^10,16^. Overton et al. concluded that p21 plays crucial role in maintaining the population heterogeneity for the MCF10A cells in a culture medium^10^. We took the same network (Fig. 1 and Fig. S1), model equations (Table S1) and modified few of our model parameters (Table S4, rest of the parameters are same as in Table S3) to obtain the E2F steady state bifurcation diagram as a function of GF for the WT (Fig. 5a, WT) and p21 deficient case (Fig. 5a, p21^-/-^).

**Fig. 5.**
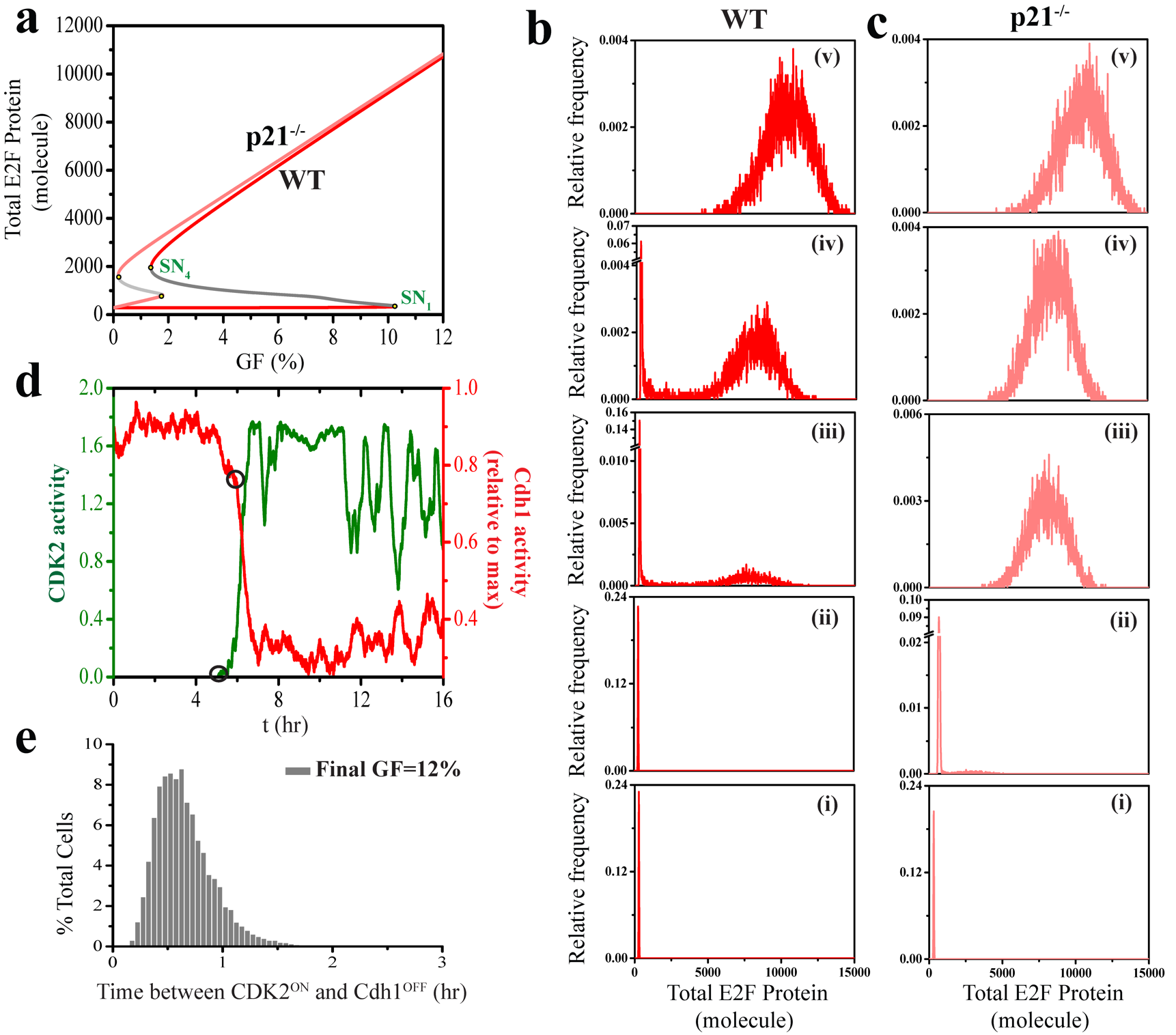
p21 governs the growth factor-dependent population heterogeneity in MCF10A cells. **(a)** The E2F steady state dynamics as a function of GF displays an extended bi-stable domain in wild type situation (**WT**), whereas the bi-stable region reduces drastically under p21 deficient (**p21**^-/-^) condition. Numerically simulated histograms of total E2F level are shown for 10000 cells (i.e., in our case for 10000 different starting random number seeds during stochastic simulation) after 48 hrs of non-pulsed GF stimulation for **(b) WT** (red line) and **(c) p21**^-/-^ (shaded red line) situations. Final GF percentages are set to **(i)** 0.1% **(ii)** 1.5% **(iii)** 8.5%, **(iv)** 9%, and **(v)** 12%, respectively. **(d)** Representative stochastic trajectories for Cdh1 (red line) and CDK2 (green line) activities under non-pulsed GF stimulation (GF = 12%) show that irreversible Cdh1 inactivation sets in after the rise of CDK2 activity. **(e)** Histogram of the time between CDK2^ON^ and Cdh1^OFF^ state (demarcated by the black circles in **(d)**) as calculated for a population of cells (considering 10000 different random number seed in stochastic simulations). Parameters adjusted to reconcile MCF10A cell-type features are mentioned in Table S4. Rest of the parameters is same as Table S3. In case of **p21**^-/-^ transcription and translation rates related to p21 synthesis are set to zero (*k*_y3_=0 molecule min^−1^, *k*_p21f_ =0 min^−1^).

Our stochastic simulation reveals that at low growth factor stimulation (like GF = 0.1% or 1.5%, Fig. 5b-c(i-ii)) both WT and p21-deficient cells remain in the E2F OFF state (quiescent G_0_ or G_1_ like state), whereas for high growth factor condition (like GF = 12%, Fig. 5b-c(v)) cells attain E2F ON state in both the cases. Importantly, in the intermediate range of growth factor stimulation (between GF = 8% to 10%) WT population distributions (Fig. 5b(iii-iv), left panel) show bi-modal E2F response, indicating cells coexisting in both E2F ON and OFF states. Thus, E2F response shows growth factor-dependent graded change in the population heterogeneity in case of WT in MCF10A. However, lack of heterogeneity was evident for the p21-deficient case under all GF conditions. These results are in line with the experimental observations made by Overton et al.^10^.

To understand the effect of Skp2 in controlling the p21-dependent population heterogeneity, we performed further stochastic simulations by blocking the Skp2-mediated degradation of p21 (Fig. S5a), and by blocking the proteasomal degradation of p21 along with Skp2-mediated degradation (Fig. S5b) with a strong growth factor pulse (GF=12%). Fig. S5a shows similar population distribution patterns as observed in case of control (Fig. 5b, WT, GF=12%), whereas Fig. S5b exhibits quiescent state (E2F OFF state) attainment at the same growth factor stimulation. These essentially support the observation made by Overton et al.^10^ that hindrance of Skp2-mediated degradation of p21 alone cannot inhibit cell cycle activity, as it is not as active as WT p21.

In a similar note, recently, Cappell et al. showed that an irreversible APC^Cdh1^ inactivation is necessary to fully commit for cell cycle entry by taking MCF10A cells as the model system^16^. They further stressed that the inactivation of APC^Cdh1^ requires activation of CDK2^16^. To verify this, we have plotted (Fig. 5d) the dynamics of CDK2 activation and Cdh1 inactivation in a representative single cell, which shows that there is indeed a bit of time delay between the CDK2 activation and Cdh1 inactivation dynamics. We have calculated the time difference between the CDK2^ON^ and Cdh1^OFF^ state for about 10000 cells and the corresponding histogram is depicted in Fig. 5e. This qualitatively captures the experimentally observed time differences between the CDK2^ON^ and Cdh1^OFF^ states for the MCF10A cells.

### Model predicts presence of heterogeneity even under p21^-/-^ condition for MCF10A cell-type with greater control of Cyclins over E2F amplitude

To test the predictive power of the proposed model, first, we probe the lack of heterogeneity aspect of MCF10A cells under p21 deficient condition. A careful look at Fig. 2a (middle panel, showing p21^-/-^) in the Overton et al.’s experimental study^10^ suggests that even under p21 deficient situation, cycling and quiescent cells can coexist for a short range of GF condition. In our model, the p21^-/-^ condition produces a tiny bistable region in the E2F steady state. This advocates that perhaps under GF pulsed kind of experiments (as performed by Yao et al.^7^), the coexistence of committed (E2F ON state) and quiescent (E2F OFF state) cells can be observed under p21^-/-^ condition. To investigate this issue, we performed a numerical pulsed experiment under WT (Fig. 6a, green line) and p21^-/-^ (Fig. 6b, green line) cases, and compared them with the non-pulsed (Fig. 6a (WT), red line, and Fig. 6b (p21^-/-^), shaded red line) E2F distributions under same conditions. The results indeed predict (Fig. 6b(i-ii), green line) that even under p21^-/-^, coexistence of committed (E2F ON state) and quiescent (E2F OFF state) cells can be obtained for a limited range of GF, if one performs a GF pulsed (20% for 5 hrs) kind of experiment. This effect will not be visible in a non-pulsed GF increasing experiment, as correctly reported by Overton et al.^7^.

The next important question one can ask is, how the important regulators like the Cyclins (CycD and CycE) and Myc affect the amplitude and timing of quiescence to cycling phase transition in MCF10A cells. Interestingly, the bifurcation analysis (Fig. 6c) of our model foretells that simultaneous inhibition of CycD and CycE activities will affect the amplitude of E2F significantly without much altering the timing of E2F activation (position of the saddle node SN_1_ remains almost intact).

**Fig. 6.**
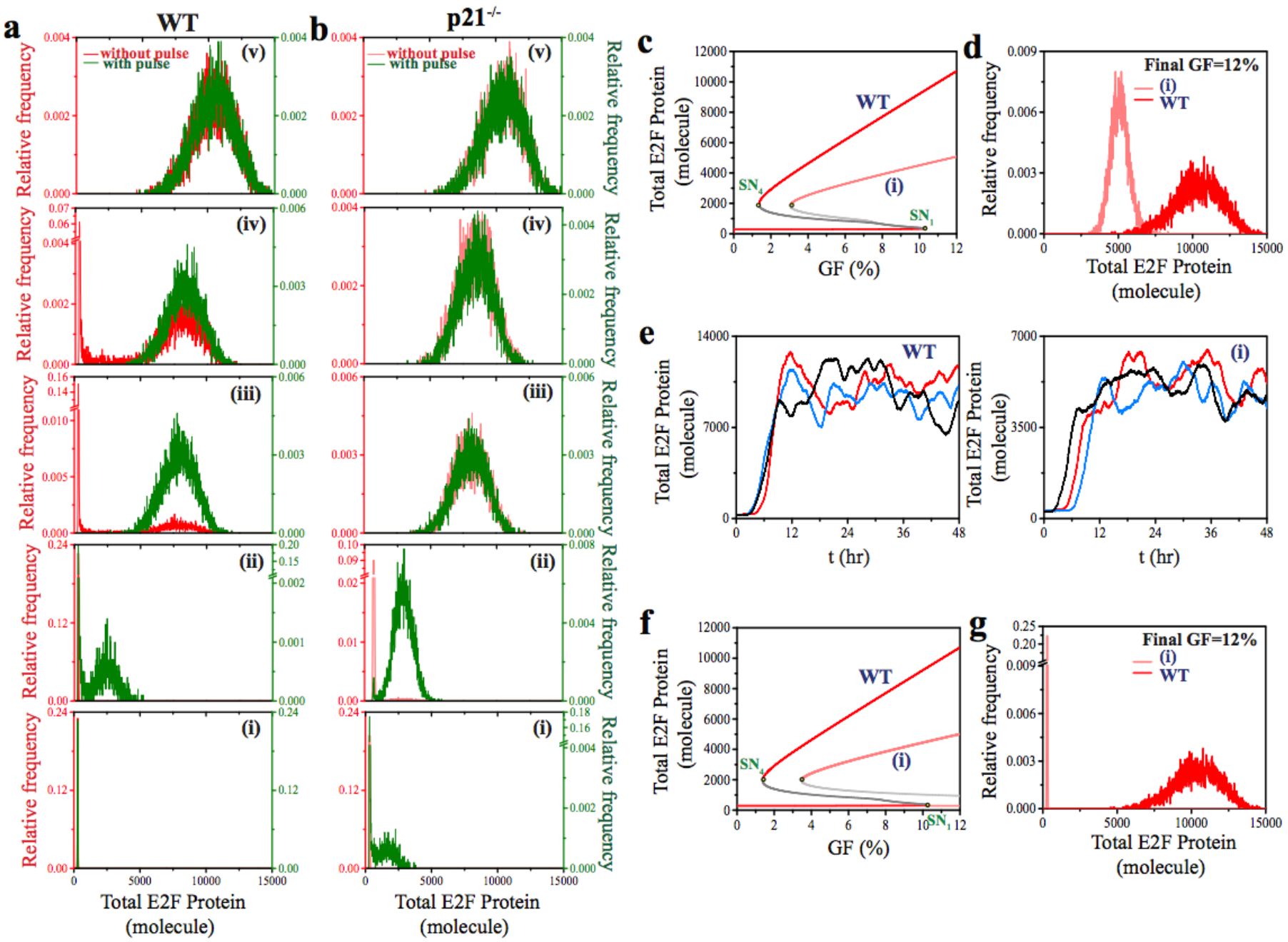
Model predicts hint of heterogeneity even under p21^-/-^ condition for MCF10A cell-type with greater control of Cyclins over E2F amplitude. Histograms of total E2F steady state for **(a) WT**, and **(b)** p21 deficient (**p21**^-/-^) condition (for 10000 cells, i.e., in our case for 10000 different starting random number seeds during stochastic simulation) after 48 hrs of GF stimulation for pulsed (20% GF for 5 hrs, green line) and non-pulsed (red line) situations. Final GF percentages are set to **(i)** 0.1% **(ii)** 1.5% **(iii)** 8.5%, **(iv)** 9%, and **(v)** 12%, respectively. **(c)** The E2F steady state dynamics as a function of GF under wild type (**WT**) and both CycD and CycE inhibitory (**(i)**, *k*_mK4_=4 molecule min^−1^, *k*_mK2_=0.4 molecule min^−1^) conditions. **(d)** The histogram (following same protocol as in Fig. 2) of E2F level for 12% GF (non-pulsed) under simultaneous CycD and CycE **(i)** inhibition (shaded red line) shows significant amplitude difference in comparison to the WT (red line) E2F steady state distribution. **(e)** Few representative stochastic trajectories of E2F activation under 12% GF (non-pulsed) condition display that duration of E2F activation in WT (**left panel**) and in the simultaneous CycD and CycE (**(i)**, *k*_mK4_=4 molecule min^−1^, *k*_mK2_=0.4 molecule min^−1^) inhibitory conditions (**right panel**) are almost identical. **(f)** The E2F steady state dynamics as a function of GF under wild type (**WT**) and for Myc inhibition (**(i)**, *k*_y9_=0.16 min^−1^) condition. **(g)** The histograms (following same protocol as in Fig. 2) of E2F level for 12% GF (non-pulsed) under normal (**WT**) and Myc inhibition **(i)** condition display that there is significant difference in the amplitude of E2F distribution under these two situations. Parameters adjusted to reconcile MCF10A cell-type features are depicted in Table S4. Rest of the parameters is same as Table S3.

Comparison of the E2F steady state distributions (Fig. 6d, at GF = 12%) obtained via stochastic simulations show a considerable difference in E2F amplitude between the WT and in presence of simultaneous CDK inhibitors. However, the representative time course trajectories reveal that the timing of E2F activation should remain unaffected under the inhibitory situation (Fig. 6e, right panel) in comparison to the WT (Fig. 6e, left panel) in case of MCF10A cells. The model predicted effect of Myc inhibition is even more intriguing. The bifurcation diagram of E2F steady state as function of GF (Fig. 6f) portrays that inhibiting Myc (**i**) will move the positions of both saddle the nodes SN_1_ and SN_4_ toward higher GF with respect to the WT scenario (**WT**). This implies that inhibiting Myc will extend the quiescent state, and will not allow cells to achieve the E2F ON state even with reasonably high GF condition. This can be nicely seen in the stochastic simulation (Fig. 6g) executed at GF = 12% under Myc inhibition situation.

We have shown that our model can be further tuned (Table S5) to reproduce the behavior of the HT29 colon cancer cells (Fig. S6) and prostate carcinoma cells (Fig. S6). Importantly, both these cell types behave like MCF10A cells as these cells are highly under the control of CKI’s (p21 and p27)^39,40^. In the SI Text, we have discussed in details how CKI’s in these cells control the transition from the quiescence to cycling state by influencing the E2F amplitude. This shows the strength and wide scope of our proposed model in delineating the G_1_-S transition propensities in a cell-type specific manner.

### Disparate feedback interactions generate cell-type specificity during G_1_-S transition

At this juncture, it is critical to understand, why our proposed model qualitatively imitates the cell-type specific proliferation commitment for different mammalian cells? More importantly, how the cell-type specificity arises in the same network controlling the G_1_-S transition, and what are the crucial factors that control this specificity? Can our model provide any further insight in this regard? To answer these key questions, we performed a systematic sensitivity analysis (taking the position of all the saddle nodes (SN_1_, SN_2_, SN_3_ and SN_4_ in Fig. 2a, and SN_1_ and SN_4_ in Fig. 5a) as the sensitivity parameters) of the proposed model again in a cell-type specific manner to investigate the relative contributions of the five modules (Fig. 7a) that mainly govern the E2F dynamics in and around the G_1_-S transition (Fig. 1 and Fig. S1). In Fig. 7b (for REF52 cells) and Fig. 7c (for MCF10A cells), we systematically vary the expression levels of the various components represented in the 5 modules described in Fig. 7a to understand their relative contributions toward quiescence maintenance and proliferation commitment.

**Fig. 7.**
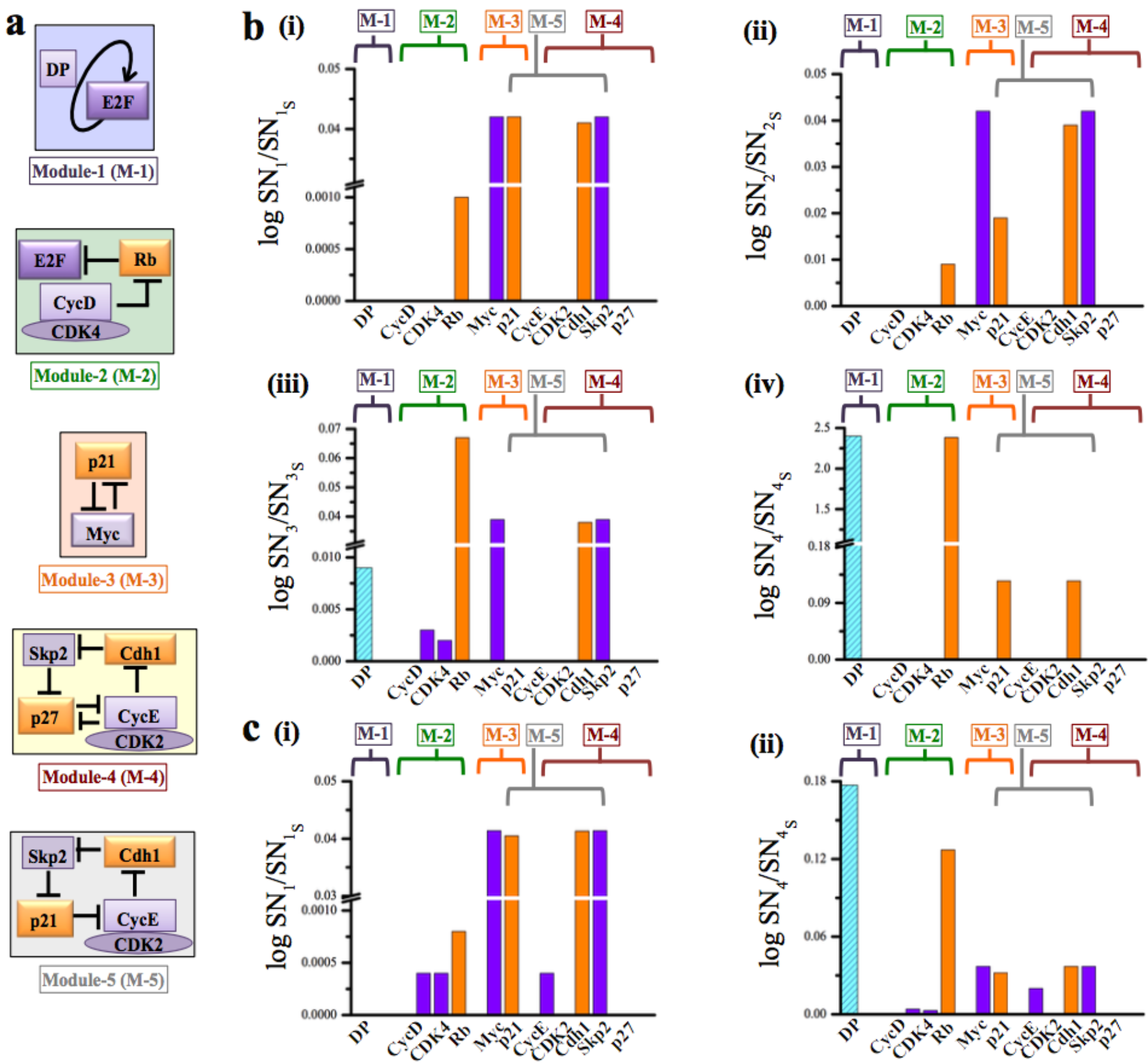
Disparate feedback interactions generate cell-type specificity during G1-S transition. **(a)** The five important modules (shown in details in Fig. 1 and Fig. S1) that dictate the cell-fate decision during G_1_-S transition of mammalian cells **(b)** Results of the sensitivity analysis performed by varying the expression levels of different network components in case of **REF52 cell-type** by taking the positions of the saddle nodes **(i)** SN_1_, **(ii)** SN_2_, **(iii)** SN_3_ and **(iv)** SN_4_ as the sensitivity parameters. **(c)** Results of the sensitivity analysis performed by varying the expression levels of different network components in case of **MCF10A cell-type** by taking the positions of the saddle nodes **(i)** SN_1_ and **(ii)** SN_2_ as the sensitivity parameters. (See the method section for details)

The sensitivity analysis reflects that the saddle nodes SN_1_ (Fig. 7b(i)) and SN_2_ (Fig. 7b(ii)), important for maintaining a quiescent G_0_ like phase in REF52 cell-type, are mainly affected by the module-3, module-5 and module-2, but the extent to which these modules alter SN_1_ and SN_2_ varies slightly. While all the three modules are responsible to determine the GF threshold (i.e. the position of SN_1_) beyond which system will transit from G_0_ like to a G_1_ like state, the extent of quiescence maintenance (determined by the difference between SN_1_ and SN_2_ in terms of GF) will mostly depend on module-3 and module-5 with a lesser dependence on module-2. However, Fig. 7b(iii) shows that the module-2 plays major role among all the modules (3,5 and 1) to decide the GF threshold for which a G_1_ like state will be maintained in REF52 cells. Fascinatingly, Fig. 7b(iv) reveals that the GF region for which a bi-stable G_1_-S transition will be maintained is mainly governed by module-1 and module-2 for this cell type. Thus, sensitivity analysis indicates that there is a clear-cut division of labor among different feedback modules to regulate the dynamics of E2F in a pre-programmed way in case of REF52 cells.

However, sensitivity analysis for the saddle node SN_1_ (Fig. 7c(i)) for the MCF10A cell type demonstrates that the GF threshold below which a quiescent like (either G_0_ or a G_1_ like state) state will be maintained is mainly governed by the module-3 and module-5 with a bit of contribution from module-2. Interestingly, Fig. 7c(i) for the MCF10A cell-type has quite a bit of resemblance with the Fig. 7b(i) obtained for REF52 cell-type with subtle important differences. In case of Fig. 7c(i), it is evident that the positioning of SN_1_ now depends on the Cyclin’s (Cyclin D/E). This shows that in case of MCF10A cells, the quiescent state has the characteristics of both G_0_ and G_1_, which was also envisaged by Overton et al.^10^. Not only that, it is further clarified that the quiescent state can now be influenced by altering the Cyclin levels for MCF10A cells, which was not possible for REF52 cells. On the other hand, the sensitivities for the saddle node SN4 delineates that almost all the modules (namely module-1, 2, 3 and 5) share the load of maintaining the bi-modal region of G_1_-S transition for the MCF10A cell type, although module-1 and 2 still contribute more relative to the other modules. While the importance of p27 related module (module-4) becomes more evident from the sensitivity analysis (Fig. S6g) performed for the HT29 colon cancer and prostate carcinoma cell-types. This clearly demonstrates that different feedback loops are operative in a dissimilar way in different mammalian cells to organize the cellular proliferation commitment in a cell type specific manner.

## Discussion and Conclusion

Proliferation commitment during late G_1_ phase of the cell cycle in mammalian cells happens in a highly cell-type dependent manner^7,9,10,40,41^. The underlying network (Fig. 1 and Fig. S1) governing the decision making events related to proliferation commitment involves several positive, negative and feed-forward kind of feedback interactions, which orchestrate the G_1_-S transition in mammalian cells by systematically influencing the dynamics of E2F. Thus, unraveling the cell-type specific distinctive nature of proliferation commitment in mammalian cells is a fascinating as well as challenging problem. In this article, our endeavor was to propose a comprehensive mathematical model of G_1_-S transition to decipher how the cell-type specific proliferation response emerges out of these complex feedback regulations operative in mammalian cells. Thus, we primarily focused our attention to two important cell-types (REF52 and MCF10A cells) for which a lot of high quality single cell experimental data is available in the literature^7,9,10,16^.

Importantly, the deterministic and stochastic analysis of our proposed model corroborate with several important experimental observation^7,9,10,16^ made for REF52 and MCF10A cells at the single cell level. Intriguingly, the model predicts for the first time that a tristable E2F steady state dynamics (Fig. 2a) in response to growth factor stimulation governs the proliferation response in rat embryonic fibroblast (REF52) cells. This not only allowed to theoretically reconcile the experimentally observed^7^ history-dependence (Fig. 2d) in the E2F dynamics, but further lead to crucial insight about the deeper quiescence maintenance (Fig. 4a-c) in the REF52 cells due to a G_0_ like state (Fig. 2a) in the E2F dynamics. At the same time, model simulation confirms the experimental observations^9,42–44^ that Myc (Fig. 3c) and Rb (Fig. 4d) critically modulate the threshold amplitude of E2F necessary to cross the R-point in REF52 cells, whereas Cyclin D/E cannot significantly alter the E2F threshold amplitude (Fig. 3a-b).

Interestingly, the model analysis reveals that for MCF10A cells, the proliferation response is organized by a bi-stable E2F steady state dynamics (Fig. 5a). In this context, our model simulations demonstrated (Fig. 5b-c) that the contribution of p21 is critical in regulating the population heterogeneity in MCF10A cells as observed by Overton et al.^10^. However, the model importantly predicts that under certain experimental condition (With GF pulse), even p21^-/-^ MCF10A cells can show population heterogeneity for a limited range of GF (Fig. 6b(i-ii)). Model further foretells the existence of history-dependence of E2F dynamics in MCF10A cells, if experiments are performed with pulsed and non-pulsed GF conditions (Fig. 6a-b). Moreover, the model analysis advocates that unlike REF52 cells, in MCF10A cells, Cyclin D/E will considerably alter the E2F threshold amplitude (Fig. 6c-d) without much affecting the time duration of E2F activation (Fig. 6e). These observations clearly display that our proposed model can be fine tuned to replicate the cell-type dependent proliferation response for both REF52 and MCF10A cell types. Not only that, we have shown that the model can be further tuned to reconcile experimental data^39,40^ for other cell-types as well.

At this point, we must highlight the fact that in literature, there were many attempts to model the G_1_-S transition for mammalian cells^17–21,23^. Those models mostly concentrate either experimental data that are related to a particular cell type, or focused more on reproducing the deterministic features of the system rather than emphasizing on the fluctuations that are important to describe single cell nature of the experimental data for different cell types. Our model is unique in this regard that it not only describes the deterministic nitty gritty of different cell types, but further explains the stochastic nature of the single cell data^7,9,10,16^ of different origin and kind, and makes novel predictions. More strikingly, the systematic sensitivity analysis (Fig. 7) allowed us to unravel how the feedback loops present in the G_1_-S transition network operate in a disproportionate manner to generate the cell type specificity in the proliferation commitment.

To summarize, our proposed model has not only corroborated various intricate single cell experimental data across different mammalian cell-types^7,9,10,16^, but also made strong and experimentally viable predictions for those cell lines. The predictions made in some cases are actually challenging the current understanding that we have related to G_1_-S transition in mammalian cells. Although at the current stage, the model is operating in a qualitative fashion, constraining the model with new and probing experimental data will definitely allow to expand the scope of the model in a more quantitative manner. We strongly believe that this model will find a wide applicability in studying a complex problem like cellular proliferation in a context dependent way, as it can be tuned to any cell type. This will definitely allow us to decipher the intriguing interplay of feedback interactions in controlling cellular proliferation, and will provide us with crucial insights about cell type specific therapeutic measures to control unwanted cellular proliferation in future.

## Methods

### Deterministic analysis

The regulatory network for G_1_-S transition of mammalian cell cycle (Fig. 1 and Fig. S1) was constructed in terms of 34 ordinary differential equations. The deterministic bifurcation analysis of the model was executed using the freely available software XPP-AUT by constructing the corresponding.ode files, and the data points generated by XPP-AUT were used to draw the bifurcation diagram (Fig. 2a, for REF52 cell-type) in Origin Lab. Parameters related to some of the modules (depicted in Fig. 1 and Fig. S1), which are adjusted to obtain the bifurcations for MCF10A cells (Fig. 5a), HT29 colon cancer cells and prostate carcinoma cells (Fig. S6) are mentioned in Table S4 and Table S5, respectively (rest of the parameters is same as depicted in Table S3 used for REF52 cell-type).

For the systematic sensitivity analysis the position of the saddle nodes in Fig. 2a, Fig. 5a and Fig. S6 are taken as the sensitivity parameter. In all the cases (Fig. 7b-c and Fig. S6g), orange bar signifies movement of the corresponding saddle node towards higher GF, and violet bar signifies the movement of the same along lower GF, when the transcriptional activation rates of the corresponding mRNA of the network components are increased by 10%. Only for the case of DP protein, the transcriptional activation rate is decreased by 10% as increasing it takes the saddle node (SN_4_ for REF52 cell-type) to unphysical concentrations, thus the cyan bar (only in case of DP) signifies the movement of the corresponding saddle nodes towards higher GF.

### Stochastic simulations

The deterministic mathematical model, constructed in terms of mass action kinetics (except one term), is transformed into a stochastic model by using the Gillespie’s stochastic simulation algorithm (SSA)^24^, which essentially takes into account the molecular fluctuation (intrinsic noise) present in a network. The reactions and the propensities of the reactions are depicted in Table S6. In order to mimic the FACS experiments performed in different experimental studies^7,9,10,16,38^, we performed our SSA simulations at each instance by changing the starting random number seed of the uniformly distributed random number generator. This allowed us to simulate 10000 individual trajectories with different random number seeds that can be compared with the data obtained in a FACS experiment comprising of ~10000 cells. We initialized the dynamical system at very low GF (say around GF = 0.0005%, which theoretically represents a quiescent state), and adopted those initial states as the initial condition for each stochastic simulation run. Corresponding histograms were plotted for 10000 cells for pulsed (20% GF for 5 hrs, green line) and non-pulsed (red line) situations after 24 or 48 hrs of simulation time depending on the experimental conditions. To compare the histograms with Yao et al.’s^7^ FACS experiment, we have expressed the x-axis of Fig. 2(d) in terms of GFP-fluorescence (where, 1 GFP-FLU = 32 molecules of E2F (case (i)), 30 molecules of E2F (case (ii)), 35 molecules of E2F (case (iii)), 38 molecules of E2F (case (iv)), 33 molecules of E2F (case (v)), 38 molecules of E2F (case (vi)), and 1 GFP-FLU = 0.6 molecules of CycD (case (i)), 3 molecules of CycD (case (ii)), 6 molecules of CycD (case (iii)), 11 molecules of CycD (case (iv)), 20 molecules of CycD (case (v)), 29 molecules of CycD (case (vi)) respectively). This has allowed us to compare our simulation output with that of the experimental data in a more comparable manner.

## Author Contribution

D.S., V.P.S.K and S.K. designed the research problem, D.S. developed the mathematical model together with V.P.S.K and S.K., D.S., V.P.S.K (partly) and S.K. analyzed the modeling outputs and predictions, D.S. and S.K. wrote the paper together.

## Acknowledgements

Thanks are due to IRCC, IIT Bombay (13IRTAPSG005) for a fellowship to (DS). We also thank UGC for providing the UGC-CSIR-JRF fellowship (Ref. No: 17- 06/2012(i)EU-V) to (VPSK). This work is supported by the funding agencies IRCC, IIT Bombay (13IRCCSG008), DST-SERB, India grant (EMR/2014/000500) and DBT, India grant (BT/PR11932/BRB/10/1315/2014).

## Conflict of Interest

The authors declare that they have no conflict of interest.

## References

(1) Hirao, A.; Arai, F.; Suda, T. Regulation of Cell Cycle in Hematopoietic Stem Cells by the Niche. Cell Cycle 2004, 3, 1481–1483.

(2) Viatour, P. Bridges between Cell Cycle Regulation and Self-Renewal Maintenance. Genes and Cancer 2012, 3 (11-12), 670–677.

(3) Burdon, T.; Smith, A.; Savatier, P. Signalling, Cell Cycle and Pluripotency in Embryonic Stem Cells. Trends Cell Biol. 2002, 12 (9), 432–438.

(4) Simons, B. D.; Clevers, H. Strategies for Homeostatic Stem Cell Self-Renewal in Adult Tissues. Cell 2011, 145 (6), 851–862.

(5) Beck, B.; Blanpain, C. Mechanisms Regulating Epidermal Stem Cells. EMBO J. 2012, 31 (9), 2067–2075.

(6) Passegué, E.; Wagers, A. J.; Giuriato, S.; Anderson, W. C.; Weissman, I. L. Global Analysis of Proliferation and Cell Cycle Gene Expression in the Regulation of Hematopoietic Stem and Progenitor Cell Fates. J. Exp. Med. 2005, 202 (11), 1599–1611.

(7) Yao, G.; Lee, T. J.; Mori, S.; Nevins, J. R.; You, L. A Bistable Rb-E2F Switch Underlies the Restriction Point. Nat. Cell Biol. 2008, 10 (4), 476–482.

(8) Aguda, B. D.; Kim, Y.; Piper-Hunter, M. G.; Friedman, A.; Marsh, C. B. MicroRNA Regulation of a Cancer Network: Consequences of the Feedback Loops Involving MiR-17-92, E2F, and Myc. Proc. Natl. Acad. Sci. 2008, 105 (50), 19678–19683.

(9) Dong, P.; Maddali, M. V.; Srimani, J. K.; Thélot, F.; Nevins, J. R.; Mathey-Prevot, B.; You, L. Division of Labour between Myc and G1 Cyclins in Cell Cycle Commitment and Pace Control. Nat. Commun. 2014, 5.

(10) Overton, K. W.; Spencer, S. L.; Noderer, W. L.; Meyer, T.; Wang, C. L. Basal P21 Controls Population Heterogeneity in Cycling and Quiescent Cell Cycle States. Proc. Natl. Acad. Sci. 2014, 111 (41), E4386–E4393.

(11) Yao, G.; Tan, C.; West, M.; Nevins, J. R.; You, L. Origin of Bistability Underlying Mammalian Cell Cycle Entry. Mol. Syst. Biol. 2011, 7 (485), 1–10.

(12) Wong, J. V.; Dong, P.; Nevins, J. R.; Mathey-Prevot, B.; You, L. Network Calisthenics: Control of E2F Dynamics in Cell Cycle Entry. Cell Cycle 2011, 10 (18), 3086–3094.

(13) Helin, K. Regulation of Cell Proliferation by the E2F Transcription Factors. Curr. Opin. Genet. Dev. 1998, 8 (1), 28–35.

(14) Helin, K.; Wu, C. L.; Fattaey, A. R.; Lees, J. A.; Dynlacht, B. D.; Ngwu, C.; Harlow, E. Heterodimerization of the Transcription Factors E2F-1 and DP-1 Leads to Cooperative Trans-Activation. Genes Dev. 1993, 7 (10), 1850–1861.

(15) Spencer, S. L.; Cappell, S. D.; Tsai, F. C.; Overton, K. W.; Wang, C. L.; Meyer, T. The Proliferation-Quiescence Decision Is Controlled by a Bifurcation in CDK2 Activity at Mitotic Exit. Cell 2013, 155 (2), 369–383.

(16) Cappell, S. D.; Chung, M.; Jaimovich, A.; Spencer, S. L.; Meyer, T. Irreversible APC^Cdh1^ Inactivation Underlies the Point of No Return for Cell-Cycle Entry. Cell 2016, 166 (1), 167–180.

(17) Barr, A. R.; Heldt, F. S.; Zhang, T.; Bakal, C.; Novák, B. A Dynamical Framework for the All-or-None G1/S Transition. Cell Syst. 2016, 2 (1), 27–37.

(18) Gerard, C.; Goldbeter, A. The Balance between Cell Cycle Arrest and Cell Proliferation: Control by the Extracellular Matrix and by Contact Inhibition. Interface Focus 2014, 4 (3), 20130075–20130075.

(19) Gerard, C.; Goldbeter, A. Temporal Self-Organization of the Cyclin/Cdk Network Driving the Mammalian Cell Cycle. Proc. Natl. Acad. Sci. 2009, 106 (51), 21643–21648.

(20) Novák, B.; Tyson, J. J. A Model for Restriction Point Control of the Mammalian Cell Cycle. J. Theor. Biol. 2004, 230, 563–579.

(21) Conradie, R.; Bruggeman, F. J.; Ciliberto, A.; Csikász-Nagy, A.; Novák, B.; Westerhoff, H. V.; Snoep, J. L. Restriction Point Control of the Mammalian Cell Cycle via the Cyclin E/Cdk2:P27 Complex. FEBS J. 2010, 277, 357–367.

(22) Verdugo, A.; Vinod, P. K.; Tyson, J. J.; Novak, B. Molecular Mechanisms Creating Bistable Switches at Cell Cycle Transitions. Open Biol. 2013, 3, 120179.

(23) Gérard, C.; Goldbeter, A. From Quiescence to Proliferation: Cdk Oscillations Drive the Mammalian Cell Cycle. Front. Physiol. 2012, 3, 1–18.

(24) Gillespie, D. T. A General Method for Numerically Simulating the Stochastic Time Evolution of Coupled Chemical Reactions. J. Comput. Phys. 1976, 22, 403–434.

(25) Chellappan, S. P.; Hiebert, S.; Mudryj, M.; Horowitz, J. M.; Nevins, J. R. The E2F Transcription Factor Is a Cellular Target for the RB Protein. Cell 1991, 65 (6), 1053–1061.

(26) Coller, H. A. From Quiescence Versus Proliferation. Mol. cell Biol. 2007, 8, 667–670.

(27) Yuan, X.; Srividhya, J.; De Luca, T.; Lee, J.-h. E.; Pomerening, J. R. Uncovering the Role of APC-Cdh1 in Generating the Dynamics of S-Phase Onset. Mol. Biol. Cell 2014, 25 (4), 441–456.

(28) Yoon, M. K.; Mitrea, D. M.; Ou, L.; Kriwacki, R. W. Cell Cycle Regulation by the Intrinsically Disordered Proteins P21 and P27. Biochem. Soc. Trans. 2012, 40, 981–988.

(29) Heldt, F. S.; Barr, A. R.; Cooper, S.; Bakal, C.; Novák, B. A Comprehensive Model for the Proliferation-Quiescence Decision in Response to Endogenous DNA Damage in Human Cells. Proc. Natl. Acad. Sci. 2018, 201715345.

(30) Lu, Z.; Hunter, T. Ubiquitylation and Proteasomal Degradation of the p21^Cip1^, p27^Kip1^ and p57^Kip2^ CDK Inhibitors. Cell Cycle 2010, 9 (12), 2342–2352.

(31) Montagnoli, A; Fiore, F.; Eytan, E.; Carrano, A. C.; Draetta, G. F.; Hershko, A.; Pagano, M. Ubiquitination of p27 Is Regulated by CDK Dependent Phosphorylation and Trimeric Comples Formation. Genes Dev 1999, 13, 1181–1189.

(32) Sheaff, R. J.; Groudine, M.; Roberts, J. M. Cyclin E-CDK2 Is a Regulator of p27^Kip1^. 1997, 1464–1478.

(33) Johnson, D. G.; Ohtani, K.; Nevins, J. R. Autoregulatory Control of E2F1 Expression in Response to Positive and Negative Regulators of Cell Cycle Progression. Genes Dev. 1994, 8 (13), 1514–1525.

(34) Leung, J. Y.; Ehmann, G. L.; Giangrande, P. H.; Nevins, J. R. A Role for Myc in Facilitating Transcription Activation by E2F1. Oncogene 2008, 27 (30), 4172–4179.

(35) Kitaura, H.; Shinshi, M.; Uchikoshi, Y.; Ono, T.; Tsurimoto, T.; Yoshikawa, H.; Iguchi-ariga, S. M. M.; Ariga, H. Reciprocal Regulation via Protein-Protein Interaction between c-Myc and p21^Cip1/Waf1/Sdi1^ in DNA Replication and Transcription. J. Biol. Chem. 2000, 275 (14), 10477–10483.

(36) Dotto, G. P. p21^WAF1/Cip1^: More than a Break to the Cell Cycle? Biochim. Biophys. Acta - Rev. Cancer 2000, 1471 (1).

(37) Mukherjee, S.; Conrad, S. E. c-Myc Suppresses p21^WAF1/CIP1^ Expression during Estrogen Signaling and Antiestrogen Resistance in Human Breast Cancer Cells. J. Biol. Chem. 2005, 280 (18), 17617–17625.

(38) Kwon, J. S.; Everetts, N. J.; Wang, X.; Wang, W.; Della Croce, K.; Xing, J.; Yao, G. Controlling Depth of Cellular Quiescence by an Rb-E2F Network Switch. Cell Rep. 2017, 20 (13), 3223–3235.

(39) Yamamoto, H.; Soh, J. W.; Shirin, H.; Xing, W. Q.; Lim, J. T.; Yao, Y.; Slosberg, E.; Tomita, N.; Schieren, I.; Weinstein, I. B. Comparative Effects of Overexpression of p27^Kip1^ and p21^Cip1/Waf1^ on Growth and Differentiation in Human Colon Carcinoma Cells. Oncogene 1999, 18 (1), 103–115.

(40) Roy, S.; Kaur, M.; Agarwal, C.; Tecklenburg, M.; Sclafani, R. A.; Agarwal, R. p21 and p27 Induction By Silibinin Is Essential for Its Cell Cycle Arrest Effect in Prostate Carcinoma Cells. Mol. Cancer Ther. 2007, 6 (10), 2696–2707.

(41) Ochiai, H.; Sugawara, T.; Sakuma, T.; Yamamoto, T. Stochastic Promoter Activation Affects Nanog Expression Variability in Mouse Embryonic Stem Cells. Sci. Rep. 2014, 4, 7125.

(42) Jacks, T.; Fazeli, A.; Schmitt, E. M.; Bronson, R. T.; Goodell, M. A.; Weinberg, R. A. Effects of an Rb Mutation in the Mouse. Nature 1992, 359 (6393), 295–300.

(43) Lee, E. Y. H. P.; Chang, C. Y.; Hu, N.; Wang, Y. C. J.; Lai, C. C.; Herrup, K.; Lee, W. H.; Bradley, A. Mice Deficient for Rb Are Nonviable and Show Defects in Neurogenesis and Haematopoiesis. Nature 1992, 359 (6393), 288–294.

(44) Sage, J.; Mulligan, G. J.; Attardi, L. D.; Miller, A.; Chen, S.; Williams, B.; Theodorou, E.; Jacks, T. Control and Immortalization Targeted Disruption of the Three Rb-Related Genes Leads to Loss of G_1_ Control and Immortalization. Genes Dev. 2000, 3037–3050.

